# Age and genetic background modify hybrid male sterility in house mice

**DOI:** 10.1101/2020.06.26.173047

**Authors:** Samuel J. Widmayer, Mary Ann Handel, David L. Aylor

**Affiliations:** Department of Biological Science, W.M. Keck Center for Behavioral Biology, and Graduate Program in Genetics, North Carolina State University, Raleigh, NC 27695; The Jackson Laboratory, Bar Harbor, ME, 04609; Bioinformatics Research Center, Center for Human Health and the Environment, and Comparative Medicine Institute, North Carolina State University, Raleigh, NC 27695

**Keywords:** Hybrid Male Sterility, Aging, Genetic Background, Spermatogenesis, Mouse

## Abstract

Hybrid male sterility (HMS) contributes to reproductive isolation commonly observed among house mouse *(Mus musculus)* subspecies, both in the wild and in laboratory crosses. Incompatibilities involving specific *Prdm9* alleles and certain Chromosome (Chr) X genotypes are known determinants of fertility and HMS, and previous work in the field has demonstrated that genetic background modifies these two major loci. We constructed hybrids that have identical genotypes at *Prdm9* and identical X chromosomes, but differ widely across the rest of the genome. In each case, we crossed female PWK/PhJ mice representative of the *M. m. musculus* subspecies to males from a classical inbred strain representative of *M. m. domesticus:* 129S1/SvImJ, A/J, C57BL/6J, or DBA/2J. We detected three distinct trajectories of fertility among the hybrids using breeding experiments. The PWK129S1 males were always infertile. PWKDBA2 males were fertile, despite their genotypes at the major HMS loci. We also observed age-dependent changes in fertility parameters across multiple genetic backgrounds. The PWKB6 and PWKAJ males were always infertile before 15 weeks and after 35 weeks, yet some PWKB6 and PWKAJ males were fertile between fifteen and 35 weeks. This observation could resolve previous contradictory reports about the fertility of PWKB6. Taken together, these results point to multiple segregating HMS modifier alleles, some of which have age-related modes of action. The ultimate identification of these alleles and their age-related mechanisms will advance understanding both of the genetic architecture of HMS and of how reproductive barriers are maintained between house mouse subspecies.

## Introduction

Hybrid male sterility (HMS) is the phenomenon in which matings between genetically distinct parents produce viable yet infertile male offspring. It is an important contributor to reproductive isolation between populations and incipient species. The Dobzhansky-Muller model (Dobzhansky 1937; Muller 1942; Orr 1995) proposes an evolutionary genetic mechanism for the development of reproductive incompatibilities. With restriction to gene flow, diverging populations accumulate and fix new mutations. These derived alleles are neutral within each population, but act deleteriously in hybrids through epistatic interactions that cause HMS. HMS is an example of Haldane’s rule, the observation that when one sex is absent or sterile in interspecies cross progeny, that sex is the heterogametic sex (Haldane 1922). An important extension of Haldane’s rule is that, sometimes, reciprocal interspecies crosses produce sterile hybrid offspring in one cross direction, but not the other, if incompatibilities evolve at different rates in each species (Turelli and Moyle 2007).

House mice (*Mus musculus)* are a powerful system for studying HMS. House mice have a cosmopolitan distribution and exist in three genetically distinct subspecies: *M. m. musculus, M. m. domesticus,* and *M. m. castaneus* (Boursot *et al.* 1993, 1996; Phifer-Rixey and Nachman 2015). These subspecies began diverging approximately 500 thousand years ago (Geraldes *et al.* 2008), yet gene flow between the *M. m. musculus* and *M. m. domesticus* subspecies still occurs in a narrow hybrid zone in central Europe. Here the two subspecies exhibit partial reproductive isolation, with reduced fertility largely restricted to males with relatively high degrees of subspecies admixture (Turner *et al.* 2012). Subspecies-diagnostic allele frequencies exhibit sharp clines across the hybrid zone (Tucker *et al.* 1992; Teeter *et al.* 2008; Janousek *et al.* 2012). Furthermore, the degree of subspecies introgression genome-wide is heterogeneous, with a particularly strong reduction in sex chromosome introgression (Payseur *et al.* 2004). Therefore, while certain regions of the genome flow freely between subspecies, reproductive isolation is partial and maintained primarily by selection against infertile hybrid males in nature.

Studies of natural mouse populations have revealed few candidate HMS loci, and others have made progress by crossing inbred mouse strains representative of the major mouse subspecies (Gregorová and Forejt 2000; Britton-Davidian *et al.* 2005; Good *et al.* 2008b; Pialek *et al.* 2008; White *et al.* 2011; Larson *et al.* 2018). Hybrid sterility is generally asymmetric in crosses between primarily *M. m. domesticus-derived* and *M. m. musculus-derived* inbred strains, affecting only hybrid males derived from *M. m. musculus* dams and *M. m. domesticus* sires (Good *et al.* 2008b; Mihola *et al.* 2009; Flachs *et al.* 2012; Bhattacharyya *et al.* 2013).

Allelic variation at the gene *Prdm9* and the Chr X QTL *Hstx2* have been shown to play a key role in most genetic studies of mouse HMS. Specific genotypes at these HMS loci have been associated with infertility phenotypes characterized by meiotic prophase abnormalities including asynapsis during the pachytene stage (Mihola *et al.* 2009; Flachs *et al.* 2012, 2014) and impaired meiotic sex chromosome inactivation (MSCI) (Good *et al.* 2010; Bhattacharyya *et al.* 2013, 2014; Campbell *et al.* 2013; Larson *et al.* 2017). The *Prdm9* gene encodes a histone methyltransferase that is required for selection and activation of genomic recombination hotspots, a process that ultimately leads to creation of meiotic DNA double-strand breaks (DSBs) and formation of the synaptonemal complex that facilitates homologous chromosome interactions (Hayashi *et al.* 2005; Mihola *et al.* 2009; Flachs *et al.* 2012, 2014; Bhattacharyya *et al.* 2013; Davies *et al.* 2016). In fertile male mice, PRDM9 binds to specific DNA sequences in spermatocytes, thus selecting and demarcating recombination hotspots. This ultimately leads to the formation of DSBs that initiate recombination immediately preceding synapsis of homologous chromosomes (Baudat *et al.* 2010a; Parvanov *et al.* 2010). The protein isoforms of PRDM9 exhibit allelespecific binding genome-wide (Brick *et al.* 2012; Baker *et al.* 2014, 2015; Walker *et al.* 2015) and the *Prdm9^Dom2^* allele exhibits DNA-binding motif variation relative to *Prdm9^Msc^* or *Prdm9^Dom3^.* Aberrant DNA-binding in *Prdm9* heterozygotes is also associated with asynapsis (Davies *et al.* 2016). Fertility in this HMS system can be rescued by decreasing the extent of asymmetric PRDM9 binding through transgenic rescue (Mihola *et al.* 2009), replacing the *Dom2* allele with *Dom3* or humanized *Prdm9* alleles (Flachs *et al.* 2012, 2014; Davies *et al.* 2016), or by artificially creating random stretches of symmetric homology between certain chromosome pairs (Gregorova *et al.* 2018). The *M. m. musculus* allele of *Hstx2 (*Dzur-Gejdosova *et al. 2012;* Bhattacharyya *et al. 2014;* Balcova *et al*. 2016; Lustyk *et al*. 2019) is necessary for HMS in all of the reported *M. m. musculus* x *M. m. domesticus* hybrids, though the nature of its interaction with *Prdm9* remains uncharacterized.

Although *Prdm9* and *Hstx2* are important drivers of HMS, studies of genetic background effects indicate these allele incompatibilities are not sufficient to cause HMS. Wild mice captured from the hybrid zone are frequently fertile (Turner *et al.* 2012; Turner and Harr 2014) 2012), indicating that specific genetic architectures can rescue fertility. Indeed, association mapping has revealed large-effect QTL in the wild (Turner and Harr 2014). Furthermore, many alleles of *Prdm9* segregate within natural populations (Buard *et al.* 2014, Kono et al. 2014), and many are uncharacterized with respect to fertility in hybrid males. Two alleles are well-studied and also segregate within the primarily *M. m. domesticus-derived* inbred strains. Intersubspecific hybrid male mice that carry *Prdm9^Dom2^* (e.g. C57BL/6J) are generally infertile (Mihola *et al.* 2009; Flachs *et al.* 2012, 2014), while hybrid male mice that carry *Prdm9^Dom3^* (e.g, WSB/EiJ) are typically fertile in spite of severely reduced sperm count and altered sperm morphology (Good *et al.* 2008a, 2008b; Turner *et al.* 2014). Nonetheless, in F2 intercrosses derived from WSB, reproductive phenotypes segregated, and several QTL have been associated independent of *Prdm9* or Chr X (White *et al.* 2011; Turner and Harr 2014; Turner *et al.* 2014). Moreover, male progeny of C57BL/6J-Chr 17^PWD^ consomic sires and C57BL/6J-Chr x^PWD^ consomic dams carry the known HMS genotypes but are fertile, indicating the existence of additional alleles that can rescue fertility in the B6 background (Dzur-Gejdosova *et al.* 2012). HMS modifier alleles also segregate among closely related *M. m. musculus*-derived inbred mouse strains (Good *et al.* 2008a; Larson *et al.* 2018).

Such segregating modifiers are particularly relevant to observations of infertility in the Collaborative Cross (CC) (Churchill et al. 2004, 2012), a panel of mouse recombinant inbred lines that was derived from eight ancestral strains including representatives from each subspecies. A majority of incipient CC lines were lost (Shorter *et al.* 2017), presumably due to hybrid incompatibilities. However, only one of 64 possible F1 hybrids (PWK129S1) exhibited HMS (Chesler *et al.* 2008), and several of the ancestral strains shared genetic identity at *Prdm9* and *Hstx2*. Thus, while allelic variation at *Prdm9* and *Hstx2* can explain a large proportion of variation in HMS phenotypes, modifiers were clearly important to extinction of CC lines. The complete genetic architecture of HMS is not yet fully understood.

We investigated the effect of genetic background on fertility of hybrid male mice. Our strategy was to create hybrid male mice from crosses of females of the PWK/PhJ strain carrying the *Prdm9^Msc^* allele, to four different inbred mouse strains, each carrying the *Prdm9^Dom2^* allele, thus keeping both Chr X and heterozygous *Prdm9* genotype constant across the different hybrids. We measured reproductive phenotypes across the reproductive life span from 8 to 35 weeks and identified genetic background effects on age-dependent patterns of fertility. Additionally, we used public whole genome sequence data to identify genomic regions with segregating subspecific ancestry that are candidates to harbor HMS alleles. We demonstrate that undiscovered HMS alleles segregate among these strains and present a novel observation of age-dependent HMS in the mouse.

## Materials and Methods

### Mice

We generated F1 hybrid male mice by crossing PWK/PhJ (PWK) mice representative of *M. m. musculus* to four classical inbred strains: 129S1/SvImJ (129S1), A/J (AJ), C57BL/6J (B6), and DBA/2J (DBA2) that are primarily *M. m. domesticus*. All mice are referred to with the standard nomenclature of Dam Strain followed by Sire Strain (e.g. PWK129S1 males are produced by crossing PWK females to 129S1 males). MD hybrids (PWK129S1, PWKB6, PWKAJ, PWKDBA2) share *Prdm9^Dom2/Msc^* genotypes (Parvanov *et al.* 2010) and are hemizygous for the PWK Chr X. We also produced the reciprocal DM hybrids (129S1PWK, AJPWK, B6PWK, and DBA2PWK) by crossing classical inbred strain females to PWK males. These mice are also heterozygous *Prdm9^Dom2/Msc^* but vary on Chr X. All mice were fed soy-free Teklad mouse chow *ad-libitum.* All procedures involving animals were performed according to the Guide for the Care and Use of Laboratory Animals with approval by the Institutional Animal Care and Use Committee of North Carolina State University (NCSU) or the University of North Carolina at Chapel Hill (UNC-CH).

### Reproductive Phenotyping

Males were euthanized by carbon dioxide asphyxiation followed by cervical dislocation. Weights for the carcass, testes, epididymides, and seminal vesicles were recorded. The right caudal epididymis was incubated in 500 μL of phosphate-buffered saline for at least 15 minutes at 37° C in an empty petri dish. Following incubation, the vas deferens and caput epididymis were removed and the cauda was snipped and incubated again for 15 minutes at 37° C. After the second incubation, sperm were extruded from the cauda using curved forceps. Once the sperm suspension was collected in a microcentrifuge tube, the petri dish was rinsed with additional PBS to collect remaining suspension, bringing the final suspension volume to 1 mL. The sperm count was determined using a NucleoCounter SP-100 sperm cell counter (Chemometec).

### Histological Analysis

The left testis of each male was fixed in Bouin’s solution overnight, and serially washed in 25%, 50%, and 70% ethanol. Testes were then embedded in paraffin wax, sectioned at 5 μm thickness, and stained by either standard hematoxylin and eosin or periodic-acid Schiff staining protocols. From each male, 35-50 seminiferous tubule cross-sections were examined to determine frequency of meiotic and post-meiotic stages of spermatogenesis. We counted the tubules at seminiferous epithelium cycle stages III through VIII in order to capture the assessed germ-cell stages: 1) pachytene spermatocytes, 2) round spermatids, and 3) condensing spermatids.

### Fertility Testing

We conducted several independent breeding experiments in order to directly assess the fertility of test hybrids at various ages and over time. In order to determine the onset of fertility, we crossed PWKB6 (*n*=26) males around the onset of fertility at 5-8 weeks to FVB females in trio matings. Males had continuous access to females until discovery of a litter, or until euthanasia and reproductive phenotyping. We collected terminal phenotypes of nine PWKB6 males at 15-weeks old and the remaining seventeen at 20-weeks old. We included a small number of PWKAJ (*n*=6) males with the expectation that they would be similar to PWKB6. Lastly, we confirmed the fertility of the reciprocal B6PWK (*n*=6) males and PWKDBA2 (*n*=3) in the age range of 5-20 weeks.

In addition, we tested a subset of 35-week old mice at NC State to confirm their fertility status. 35-week old mice tested were as follows: AJPWK (*n*=3), B6PWK (*n*=2), DBA2PWK (*n*=3), 129PWK (*n*=3), PWKAJ (*n*=5), PWKB6 (*n*=5), PWKDBA2 (*n*=7), PWK129S1 (*n*=3). These mice were crossed with fertile FVB/NJ females at roughly 34 - weeks old and the females were monitored for litters for 22 days after being separated.

In a separate experiment conducted at UNC-CH, we tested PWKB6 male fertility after 15 weeks. PWKB6 male mice (*n*=63) were continuously crossed to FVB females beginning at ages between 18-27 weeks. Crosses were continually checked for litters until males ceased producing litters for at least four weeks. The age at which the male exhibited fertility was calculated by subtracting 21 days from the birth date of his offspring.

### Analysis of Subspecific Ancestry

Subspecific ancestry tracks for the 129S1, A/J, B6, DBA2, and PWK genomes were obtained from the publicly available data in the Mouse Phylogeny Viewer (http://msub.csbio.unc.edu/) (Wang *et al.* 2012; Yang *et al.* 2012). Each of the inbred genomes was scanned for shared subspecific ancestry with PWK using the *Granges* package implemented through Bioconductor (Lawrence *et al.* 2013). Shared genomic regions were then classified as to whether they were unique to an inbred strain (i.e. only 129S1 and PWK share common ancestry), or whether they shared subspecific ancestry with strains that, when crossed to PWK females, produce infertile hybrids at 8 weeks of age (129S1, A/J, and B6), fertile hybrids at 20 weeks of age (A/J, B6, DBA2), or hybrids that display age-dependent fertility with incomplete penetrance (A/J and B6). These regions and previously identified QTL were visualized using the BioCircos package (Krzywinski *et al.* 2009) implemented in R (version 3. 4. 4). Genomic regions that shared PWK subspecific ancestry were then queried for known genes using the UCSC mouse genome table browser. We then queried these gene sets in the Mouse Genome Informatics database (Bult *et al.* 2019) for genes previously implicated in male reproductive system function.

### Statistical Analyses

We compared MD hybrids and their corresponding reciprocals using 2-way Analysis of Variance (ANOVA) and Tukey Honestly Significant Difference (HSD) test. We compared a specific MD hybrid and its reciprocal using Welch’s t-test, which accounts for unequal sample variances. We compared the four MD hybrids using 1-way ANOVA and HSD. We compared a specific MD hybrid and its reciprocal across ages using 2-way ANOVA and HSD. We compared a specific hybrid across ages using a 1-way ANOVA and HSD. We compared multiple MD hybrids across ages using 2-way ANOVA and HSD. We performed all statistical tests using R software.

### Data Availability

Data is available on the GSA figshare portal. Table S3 contains phenotype and fertility testing data from mice bred at NCSU. Table S4 contains PWKB6 fertility testing data from a separate experiment conducted at UNC-CH.

## Results

### Genetic differences among classical inbred mouse strains shape variation in HMS phenotypes

We crossed PWK mice with mice from four different classical inbred mouse strains: 129S1, A/J, B6, and DBA2. These crosses yielded eight distinct types of hybrid male mice. All eight types were identically heterozygous at the *Prdm9* gene. The four MD hybrid males were offspring of a PWK dam, and carried the *M. m. musculus* Chr X that has previously been associated with HMS in mice (**Figure 1A**). The reciprocal DM hybrid males were offspring of a PWK sire, and carried Chr X from a classical inbred strain (**Figure 1B**). Since the MD hybrid males had invariant *Prdm9* and Chr X genotypes, observed variation in HMS traits among them was necessarily due to other genetic background features. We measured testes weight and epididymal sperm counts from replicates of each hybrid at three ages: 8 weeks, 20 weeks, and 35 weeks.

**Figure 1.**
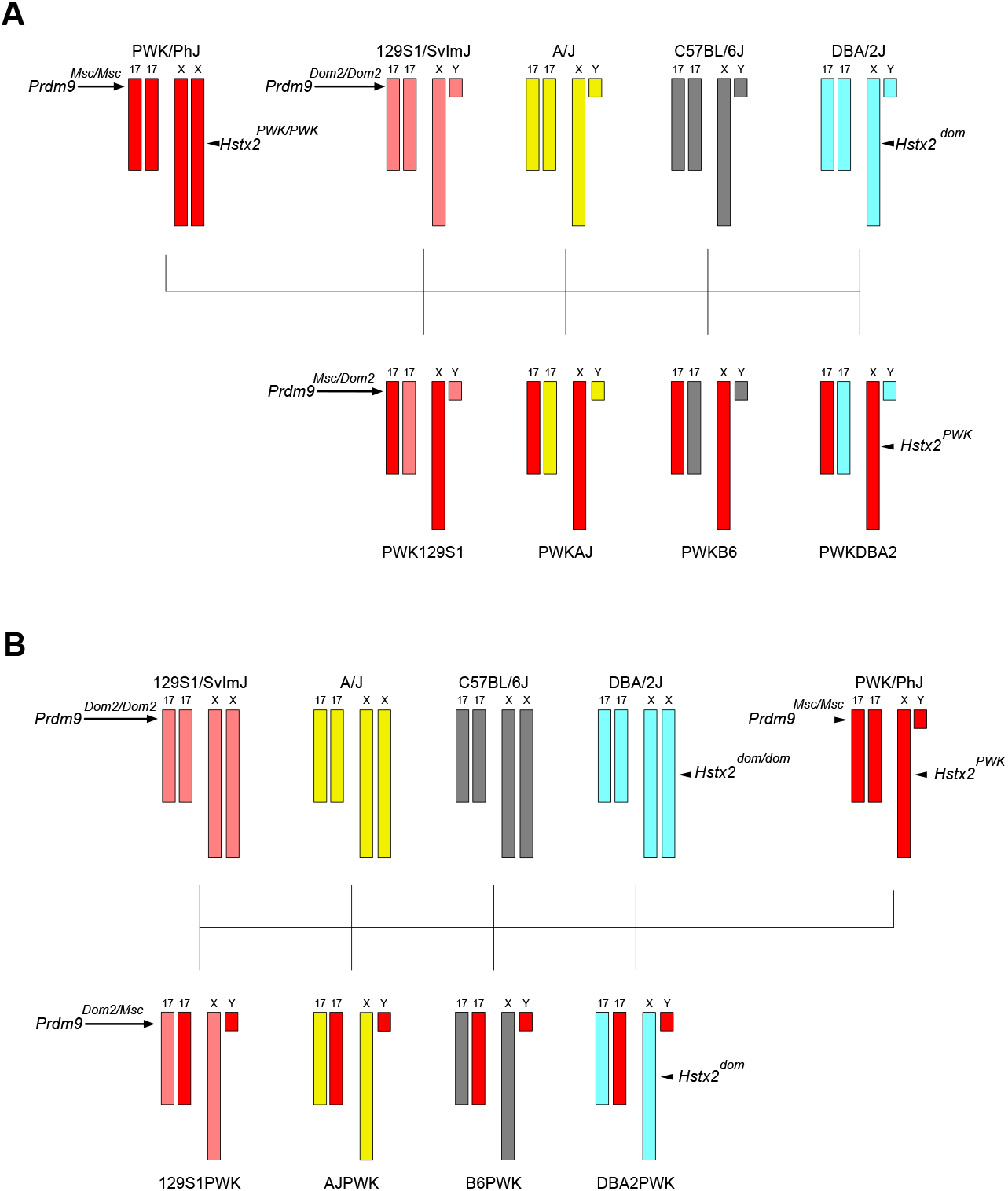
Crossing schemes to generate genetically diverse hybrid male mice. A) MD hybrid males were generated by crossing PWK (red) females to males of four classical inbred mouse strains: 129S1 (pink), A/J (yellow), B6 (gray), and DBA2 (light blue). These four strains each carry the *Prdm9^Dom2^* allele that has been previously linked to HMS. As illustrated here, MD mice are fixed for the *Prdm9^Dom2/Msc^* genotype and PWK Chr X, eliminating the effects of those major HMS loci. B) The reciprocal DM hybrids were generated by reversing the cross direction, such that Chr X is inherited from the classical inbred strain instead of PWK.

The PWK129S1, PWKAJ, and PWKB6 hybrid males (MD) had reduced combined testes weights (**Figure 2A**) and total sperm counts (**Figure 2B**) relative to their reciprocal (DM) hybrids at each age (two-way ANOVA, *p* ≤ 0.001, **Table S1**). In contrast, the PWKDBA2 (MD) testes weights were indistinguishable from reciprocal DBA2PWK (DM) males at age 8 weeks (t-test, *p* = 0.782), although PWKDBA2 testes weights were less at 20 weeks and 35 weeks (t-tests, *p* ≤ 0.005). The PWKDBA2 total sperm counts were lower than that of reciprocal hybrids at all ages (two-way ANOVA, *p_adj_* ≤ 0.001). Nonetheless, the PWKDBA2 males consistently displayed quantitative reproductive parameters that were greater than the other MD hybrids (one-way ANOVA w/ HSD, *p_adj_* ≤ 0.001) but less than the DM reciprocal hybrids.

**Figure 2.**
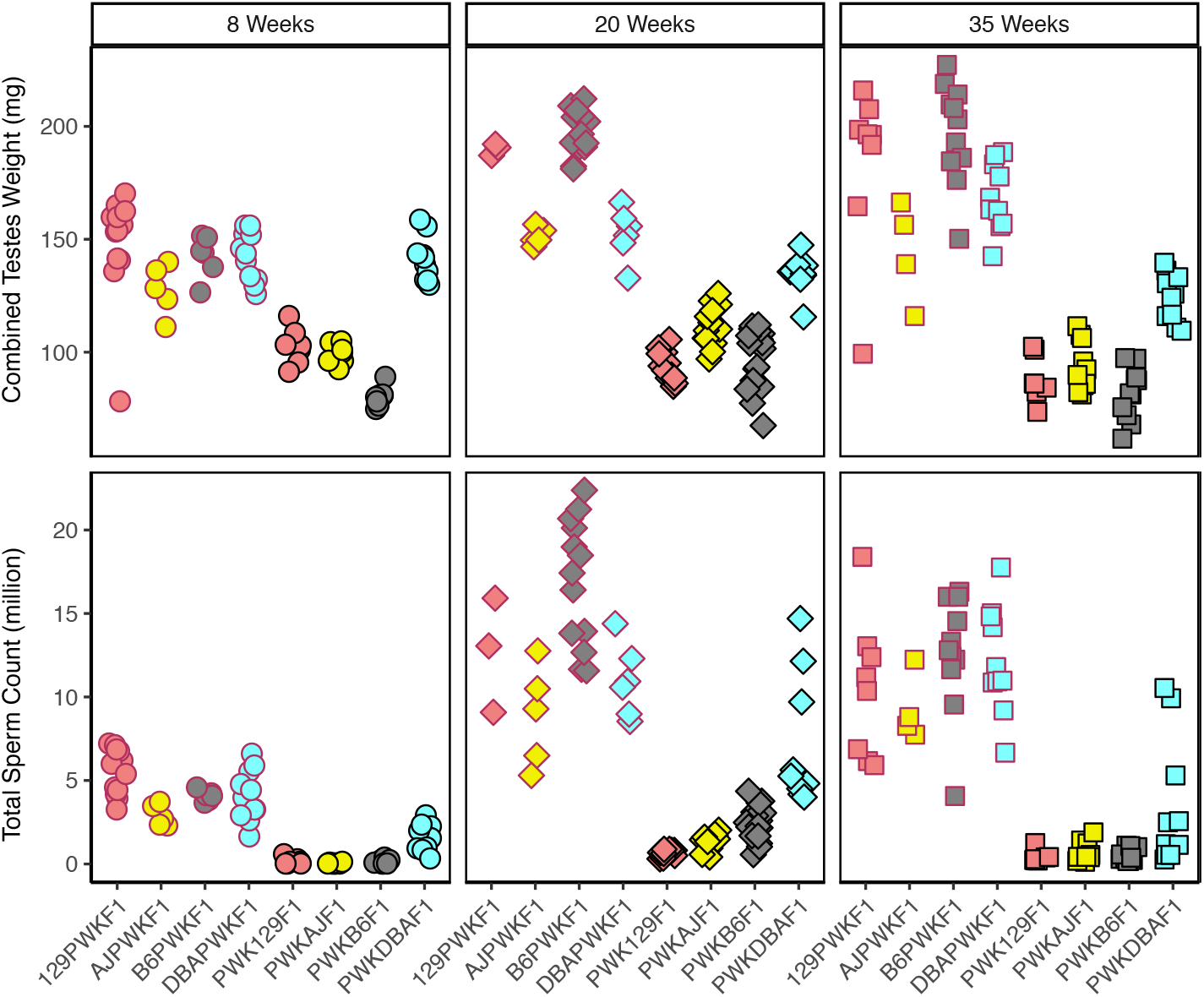
Reproductive phenotypes of hybrid male mice vary with genetic background and age. A) Combined testes weight and B) sperm count of the eight distinct hybrids across three ages: 8 weeks (circles), 20 weeks (diamonds), and 35 weeks (squares). The DM hybrid males (red borders) exhibited higher phenotypes relative to all MD hybrid males (black borders) and were fertile at all ages. PWKDBA2 (light blue with black border) exhibited higher reproductive parameters than other MD hybrids at all ages, consistent with their fertility. Slight increases in testes weights and average sperm count in PWKAJ (yellow with black border) and PWKB6 (grey with black border) hybrid males coincided with transient fertility in those types at 20 weeks of age, which subsided by 35 weeks of age.

PWK129S1 hybrids were reported to be infertile in early stages of the Collaborative Cross (CC) breeding program (Chesler *et al.* 2008); therefore the extremely low sperm counts observed (0-1.25 million) were expected. However, the low testes weights and sperm counts of PWKB6 and PWKAJ males posed a conundrum, because both of these MD hybrids have been reported to be fertile. In fact, fertile PWKAJ males and fertile PWKB6 males were necessary contributors to both the CC and the Diversity Outbred (DO) mouse resources (Churchill *et al.* 2004, 2012; Collaborative Cross Consortium 2012). Nevertheless, we observed no difference in 8-week sperm count relative to the infertile PWK129S1 males (one-way ANOVA w/ HSD, *p_adj_* > 0.955). Furthermore, PWKB6 males exhibited the lowest testes weights of all the hybrids (**Table S1**), and these weights were significantly lower than both PWKAJ males (one-way ANOVA w/ HSD, *p_adj_* ≤ 0.001) and PWK129S1 males (one-way ANOVA w/ HSD, *p_adj_* ≤ 0.001).

### Fertility is age-dependent in PWKB6 and PWKAJ hybrids

All eight hybrids showed changes in testes weight and sperm count with age, but analysis of age and strain combined revealed critical differences among hybrids. For example, sperm counts increased between 8 and 20 weeks for all hybrid males (two-way ANOVA, *p* ≤ 0.001), although the magnitude varied dramatically among strains. The largest changes were seen in the hybrid types that showed the highest reproductive parameter values at 8 weeks – the DM hybrids and the fertile MD PWKDBA2. Testes weights increased by 25.25% in DM hybrids on average, and these mice exhibited a 3.1-fold increase in sperm counts. Increases in testes weights were not due simply to increasing body weight, as relative testes weights also increased (Supplemental Fig 5). We conclude that age was an important factor that affected reproductive parameters in all PWK-derived hybrids, and these age-related changes were at least partially independent of genotypes at *Prdm9* and *Hstx2*.

The changes between 8 and 20 weeks in MD hybrids resulted in fertility differences across the strains, and shed light on the mechanisms of HMS modifiers in mice. The PWK129S1 males exhibited much smaller changes in testes weight and sperm count relative to other MD hybrids, and we concluded that those mice exhibit HMS phenotypes independent of age. In contrast, PWKB6 and PWKAJ mice exhibited marked increases in testes weight and sperm count at age 20 weeks in comparison to 8 weeks (two-way ANOVA, *p_adj_* ≤ 0.001). The average PWKB6 combined testes weight increased by 21.98%, and the average sperm count displayed a 20-fold increase. The average PWKAJ combined testes weight increased by 11.97%, and sperm count increased over 60-fold. There was substantial variation in sperm count at 20 weeks in both PWKB6 (561,000 to 4.36 million) and PWKAJ (418,000 to 2.02 million). These phenotypes overlapped with the lower tails of the phenotype distributions observed in fertile mice. This result provided a compelling explanation for the discrepancy between our observations of early-life sterility at eight weeks and the fertility necessary for the CC breeding program. These results were also consistent with a previous report that PWKB6 hybrid males exhibited delayed onset of fertility (Flachs *et al.* 2014). We predicted that the reproductive changes observed in PWKB6 and PWKAJ males between 8 and 20 weeks were sufficient to escape HMS, and that these changes were driven by genetic variants that segregated among B6, AJ, and 129S1.

We tested this hypothesis by directly measuring fertility in PWKB6 hybrid males. We crossed young mice (5-8 weeks) to fertile FVB females and measured latency until the first successful mating. PWKB6 males (*n*=26) bred continuously until adulthood. Mice were collected at age 15 weeks (*n*=9) or 20 weeks (*n*=17). The majority of these males were infertile, yet three of 26 PWKB6 males (11.5%) sired litters at ages ranging from 12 to 20 weeks. Each of these litters consisted of only one pup. These litter sizes were substantially reduced compared to four litters sired by reciprocal B6PWK (*n*=6) males that ranged from 5-11 pups. A smaller cohort of PWKAJ (*n*=6) males showed similar results. Two PWKAJ sired litters of two and three pups, respectively. These results demonstrated that infertility in young PWKB6 male mice had a high but incomplete penetrance, and suggested that this result extended to PWKAJ. In addition, those PWKB6 and PWKAJ males that were fertile experienced a delay in the onset of fertility with respect to reciprocal hybrids, and showed reduced fecundity based on litter sizes.

The increased values of reproductive parameters at 20 weeks of age did not persist until 35 weeks in MD hybrids (Figure 2), indicating that the effects of age were more complex than previously reported. Combined testes weights and sperm counts declined from 20 to 35 weeks of age in PWKB6, PWKAJ and PWKDBA2 hybrids (two-way ANOVA, *p* ≤ 0.001), though these declines were not observed in reciprocal hybrids (two-way ANOVA, *p* > 0.07). 35-week old PWKB6 and PWKAJ males exhibited phenotypes comparable to those of infertile PWK129S1 mice of the same age (one-way ANOVA w/ HSD, *p_adj_* > 0.446). A second breeding experiment confirmed a marked decline in fertility after 20 weeks in PWKB6 males (*n*=63). Mice were housed with fertile FVB female mice and left to breed continuously until the last male stopped producing litters. Forty percent of PWKB6 males sired offspring during the course of the experiment, producing a lower estimate of HMS penetrance than the early-life fertility experiment. Nonetheless, the majority of PWKB6 mice were infertile. Furthermore, the number of males who sired a litter decreased with age. No PWKB6 males sired offspring after 35 weeks of age (Figure 3).

**Figure 3.**
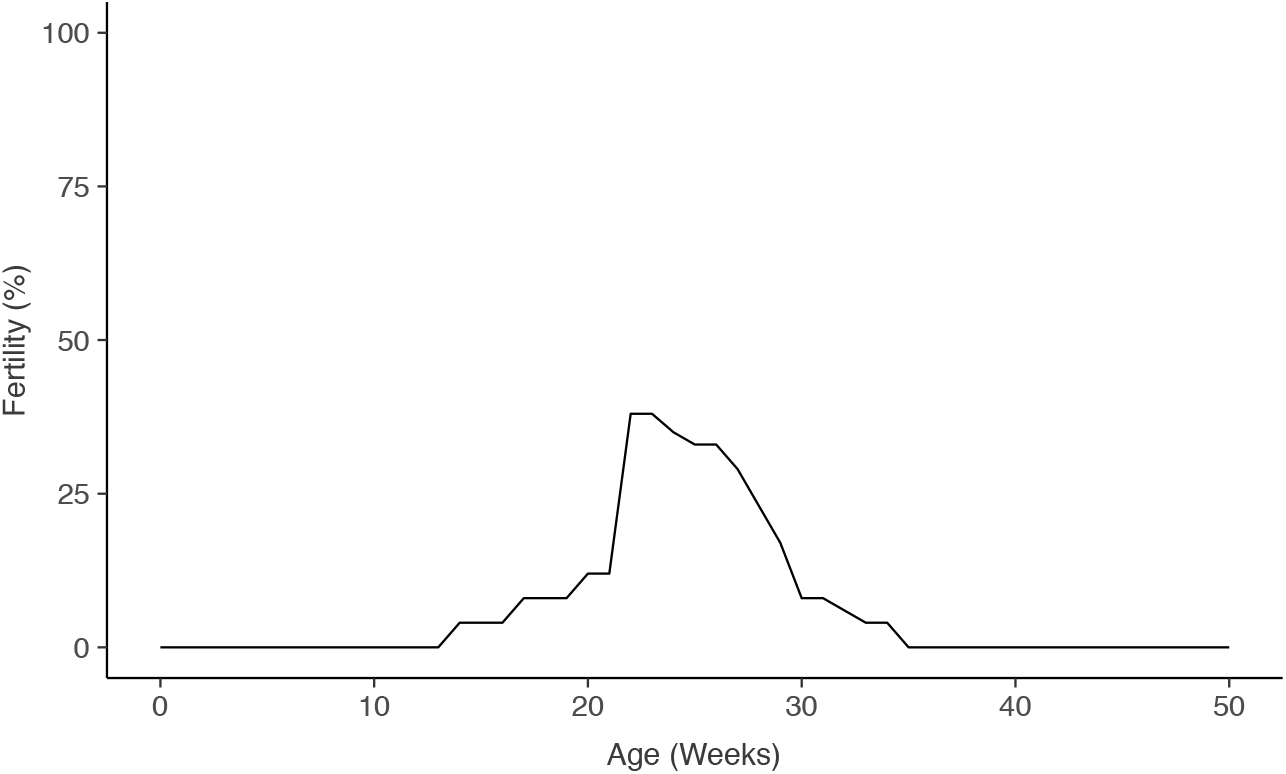
Fertility varies with age in PWKB6 hybrids. A fertility curve was constructed from combining two breeding experiments in which males had continuous access to fertile females. Most males tested were infertile. However, some mice were fertile at ages 15-35 weeks. One experiment was designed to determine the age of fertility onset and tested mice in the age range of 5-21 weeks. Three fertile males (of 26 tested) sired first litters at 15, 17, and 21 weeks. The other experiment tested fertility after age 20 weeks. 36% of males were fertile in this window, but this proportion decreased sharply with age. No males sired litters after age 35 weeks.

We concluded that age-dependent changes in testes weights and sperm count were pervasive in MD hybrid males, and that these changes were sufficient in some individuals to confer fertility during ages 12-35 weeks in PWKB6 and PWKAJ. However, this transient fertility was observed only in a minority of individuals, and we concluded that HMS was penetrant at ≥60% in these backgrounds. These results suggest at least three distinct fertility trajectories among MD hybrids: complete fertility (PWKDBA2); complete sterility (PWK129S1); and transient fertility (PWKB6, PWKAJ).

### Increased post-meiotic differentiation capacity overcomes meiotic inefficiency during the window of fertility

A defect in any stage of germ cell development could have caused quantitative sperm reductions. Multiple underlying spermatogenic impairments have previously been linked to HMS (Good *et al.* 2008b; Oka *et al.* 2010; Flachs *et al.* 2014; Schwahn *et al.* 2018). We analyzed cross-sections from hybrid male testes in order to evaluate developmental defects in sperm production (Figure 4). We focused our analysis on histological cross sections of tubules between seminiferous tubule Stages III and VIII, since the major germ cell developmental stages (spermatogonia, pachytene spermatocytes, round spermatids, and condensing spermatids) can be observed concurrently at these stages in reproductively normal mice. We measured two histological parameters. The proportion of these tubules containing round spermatids (PCT_ROUND) indicated successful spermatogenesis through completion of the meiotic division phase. The proportion of these tubules containing condensing spermatids (PCT_CONDENSING) indicated the success rate of post-meiotic differentiation from round spermatids to spermatozoa. The DM reciprocal hybrids showed virtually no evidence of spermatogenic defects, and 100% of seminiferous tubules in stages III-VIII contained both round and condensing spermatids. Similarly, the PWKDBA2 males showed no clear defects, with the exception of a slight decline in PCT_CONDENSING at age 35 weeks. Therefore, these histological analyses did not provide an obvious mechanism for how sperm count changes with age in both PWKDBA2 and all the reciprocal hybrids.

**Figure 4.**
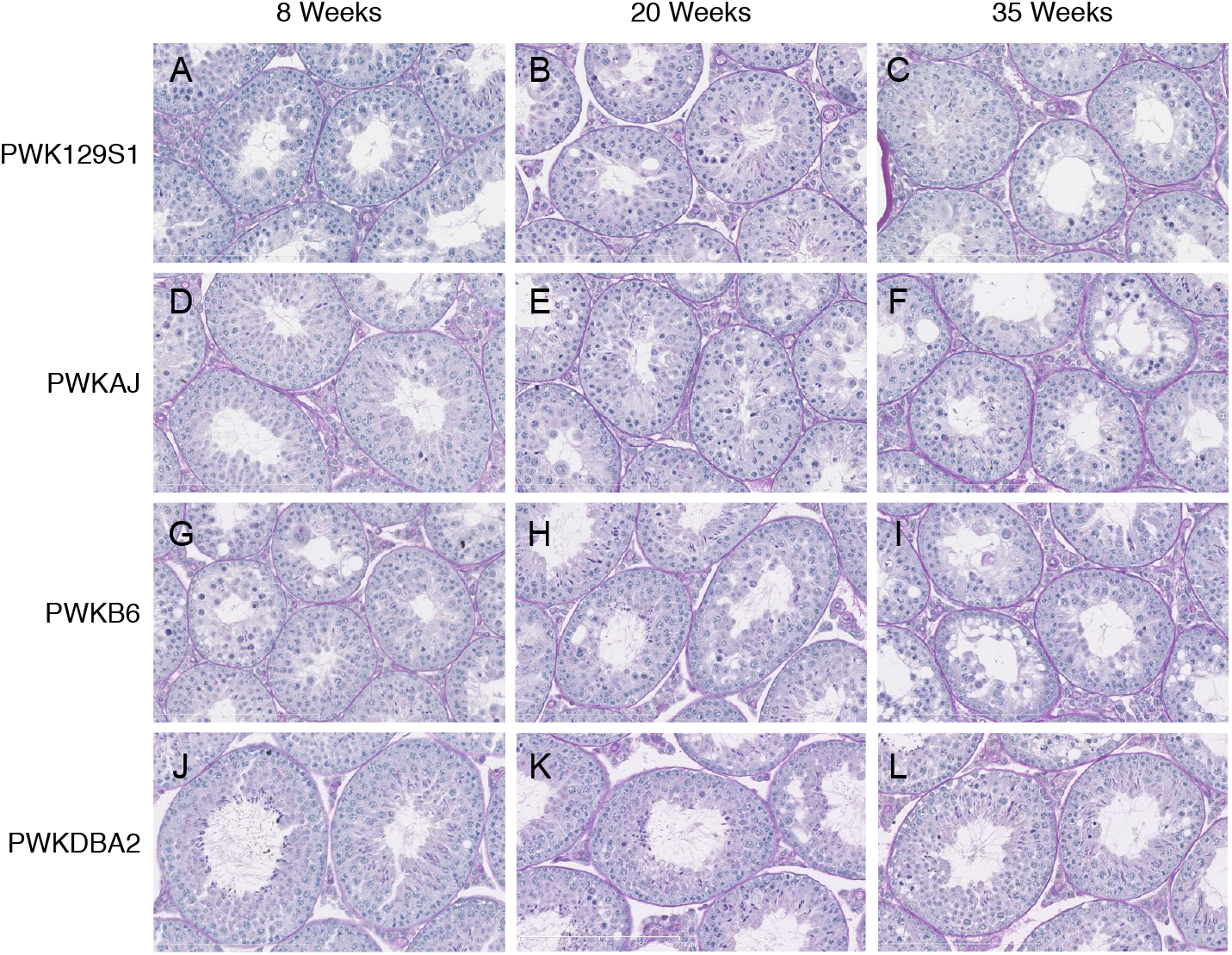
Testis histology reveals hidden complexity underlying HMS across diverse hybrid mice. Testis cross-sections stained using Periodic acid Schiff (PAS) protocol enabled staging of seminiferous tubules and analysis of distinct germ cell populations. Germ cell loss, vacuoles, and disorganization were prevalent in PWK129S1 (A-C), PWKAJ (D-F), and PWKB6 hybrids (G-I). PWKDBA2 (J-L) hybrids showed few obvious defects and were indistinguishable from DM hybrids (not shown).

The PWK129S1, PWKB6, and PWKAJ (MD) males each had a reduced PCT_ROUND relative to their reciprocal (DM) hybrids at all ages (two-way ANOVA, *p* ≤ 0.001, Figure 5A). This was consistent with their lower testes weights, total sperm counts, and fertility status. This reduction was presumably due to diminished efficiency of meiosis, consistent with previously reports of HMS associated with meiotic efficiency (White *et al.* 2011; Schwahn *et al.* 2018). We did not observe a complete arrest of meiosis, such as has been shown in other cases of HMS (Mihola *et al.* 2009; Flachs *et al.* 2012). PWK129S1 and PWKAJ males exhibited a slight age-related decrease in PCT_ROUND, and PWKB6 males exhibited no significant change with age. We concluded that meiotic efficiency explained the observed differences between two groups of hybrids: the reproductively impaired PWK129S1, PWKB6, and PWKAJ hybrids in one group, and the fertile DM and PWKDBA2 hybrids in the other. However, meiotic efficiency could not explain the transient fertility that we observed in some PWKB6 and PWKAJ hybrids.

**Figure 5.**
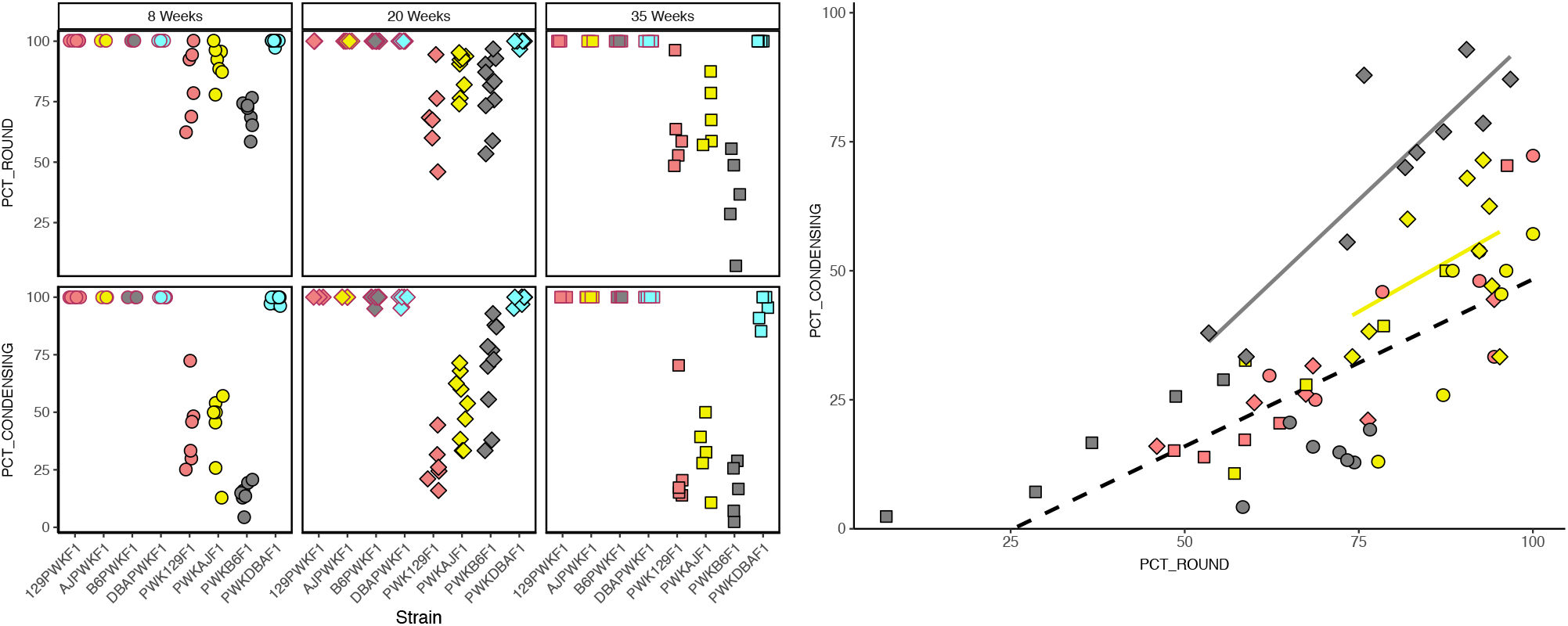
Histological parameters suggest that increased post-meiotic capacity drives age-dependent fertility. We counted the percentage of seminiferous tubules containing A) round spermatids (PCT_ROUND) and B) condensing spermatids (PCT_CONDENSING). PCT_CONDENSING increased in the transiently fertile PWKB6 (gray with black border) and PWKAJ (yellow with black border) males at 20 weeks of age. C) Since round spermatids are precursors of condensing spermatids, PCT_ROUND was correlated with PCT_CONDENSING. However, 20-week old PWKB6 (gray diamonds with black border) and PWKAJ (yellow diamonds with black border) did not show the same linear relationship as the infertile MD hybrids (all other points). The higher ratio of PCT_CONDENSING to PCT_ROUND suggests that increased post-meiotic capacity is associated with transient fertility.

In contrast, PCT_CONDENSING closely reflected the different fertility profiles that we observed among reproductively impaired hybrids. PWK129S1, PWKB6, and PWKAJ males each displayed a reduced PCT_CONDENSING compared to age-matched reciprocal hybrid males (two-way ANOVA, *p* ≤ 0.001, **Figure 5B**). This decrease could be explained simply by the reduction PCT_ROUND, since round spermatids are necessary precursors to condensing spermatids. We estimated this relationship using all the MD hybrids except for the two types that showed transient fertility – 20-week old PWKB6 and PWKAJ hybrids. There was a strong linear relationship between PCT_ROUND and PCT_CONDENSING (*r*^2^ = 0.571, **Figure 5C**). However, this relationship did not hold in 20-week old PWKAJ and PWKB6 males. These two hybrid types had substantially higher PCT_CONDENSING for a given value of PCT_ROUND than we predicted using the regression model, and also more than the infertile PWK129S1 mice of the same age (twoway ANOVA w/ HSD, *p_ad_j* ≤ 0.026). Furthermore, the marked changes with age in PCT_CONDENSING mirrored the changes in sperm count and fertility. PCT_CONDENSING increased by 55% in PWKB6 males between age 8 weeks and age 20 weeks (t-test, *p* ≤ 0.001), and increased by 10% in PWKAJ males (t-test, *p* = 0.229). This difference between PWKB6 and PWKAJ was exacerbated by the low PCT_CONDENSING value that PWKB6 males had at 8 weeks; PWKB6 males had only slightly higher PCT_CONDENSING than PWKAJ hybrids at 20 weeks (t-test, *p* = 0.052). PCT_CONDENSING declined sharply by the close of the fertility window at age 35 weeks (two-way ANOVA, p ≤ 0.001).

In summary, we conclude that two distinct processes best explained the differences observed between the eight hybrid males. The DM hybrids and the PWKDBA2 hybrids had both round and condensing spermatids in nearly all tubules, and were fertile at all ages. Reduced meiotic efficiency characterized PWKB6, PWKAJ, and PWK129S1 hybrid males, all of which were infertile at some point in their lives. However, two of this group – PWKB6 and PWKAJ – exhibited transient fertility between 15 and 35 weeks. The onset of fertility coincided with an increase in PCT_CONDENSING in our sample of 20-week old males. We interpret this to mean that the transiently fertile hybrids somehow experienced an increased capacity for the remaining round spermatids to differentiate into condensing spermatids, and this resulted in the production of enough mature sperm to confer fertility. This capacity was more robust in PWKB6 hybrids than in PWKAJ hybrids, suggesting the degree of age-dependent fertility might also vary among hybrids.

### Genomic patterns of subspecific origin implicate candidate HMS modifiers

The classical inbred strains in this study descend primarily from *M. m. domesticus* ancestors, but have important contributions from *M. m. musculus.* The surprising fertility of PWKDBA2 hybrid males could be explained if the DBA2 genome had the greatest similarity to the PWK genome, compared to the genomes of 129S1, A/J, and B6. We compared each inbred strain to PWK and isolated areas of the genome where each inbred strain shared subspecific ancestry using publicly available data (Wang *et al.* 2012). We made no assumptions about whether the mode of action of any detected incompatibilities were due to underdominance at one locus, or by acting epistatically in conjunction with at least one additional locus. In addition, we also considered loci that shared ancestry regardless of subspecies identity, since PWK also has 5.72% *M. m. domesticus* ancestry. DBA2 did not share substantially more identity overall with PWK than the other three strains. DBA2 and PWK shared subspecific ancestry across 295.7 Mb of the genome (10.83%), similar to the shared ancestry among 129S1(10.30%), B6 (9.85%), and A/J (8.39%).

Nonetheless, we reasoned that the regions of the genome shared between PWK and specific classical strains, but not others, are good candidate locations for HMS modifier alleles. We focused on four specific contrasts based on the three patterns we observed in our experiments (**Figure 6, Table S2**). First, we searched for regions of the genome where only 129S1 differed in subspecific ancestry from PWK, reasoning that these regions may be enriched for incompatibility alleles unique to the 129S1 genetic background. We identified nine such regions (7.07 Mb) across six chromosomes that contain 108 genes in total. Second, we searched for regions of the genome where A/J and B6 shared subspecific ancestry with each other but were different from DBA2 or 129S1. Regions where these strains also shared ancestry with PWK may harbor alleles that distinguish the age-dependent HMS in PWKAJ and PWKB6 males from the always-infertile PWK129S1 male. We found seven such regions (12.00 Mb) containing 146 genes. Regions in which A/J and B6 were alike but were different from the other three strains might harbor HMS alleles that explain the age-dependent effects. We found fourteen such regions (21.34 Mb) containing 111 genes. Finally, we searched for regions of the genome where only DBA2 shared subspecific ancestry with PWK. These regions might contain the critical modifier allele or alleles unique to DBA2 that rescue HMS in PWKDBA2 males. We discovered 44 such regions (87.65 Mb) containing 834 genes. These candidate regions overlap several previously identified HMS QTL (White *et al.* 2011; Dzur-Gejdosova *et al.* 2012; Bhattacharyya *et al.* 2014; Turner *et al.* 2014; Larson *et al.* 2018) (**Figure 6**) and include genes that have been previously implicated in reproductive phenotypes.

**Figure 6.**
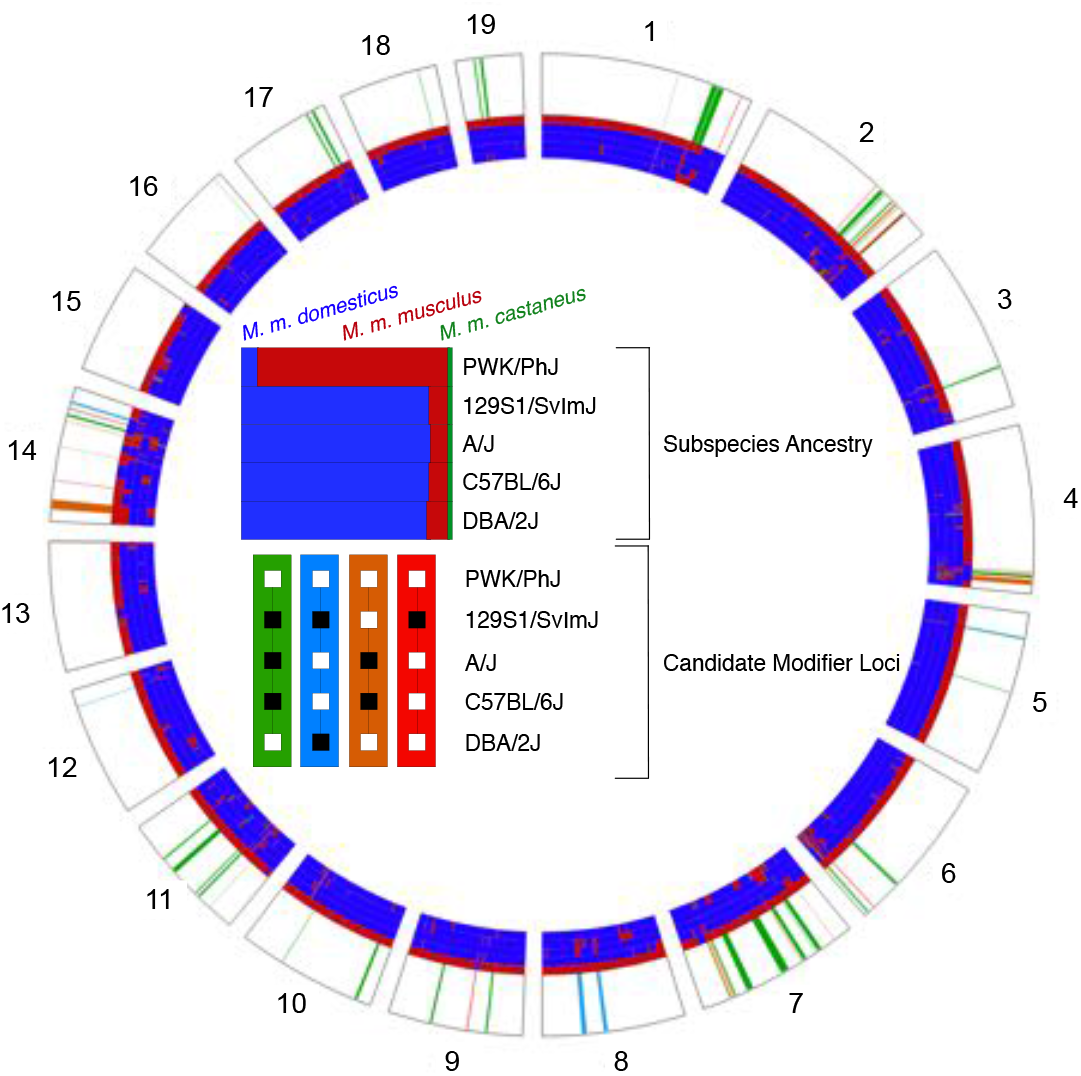
Shared ancestry reveals potential locations of HMS modifiers. Contrasting patterns of subspecific ancestry among inbred strains that contribute to the observed phenotypes. Shared regions between PWK and DBA2 (green) may harbor alleles that rescue fertility in PWKDBA2 males. Regions shared between PWK, A/J, and B6 (blue) or regions shared privately between B6 and A/J (orange) may associate with age-dependent HMS. Private incompatibilities between PWK and 129S1 (red) may underlie the complete sterility of PWK129S1 males. Gray bars indicate the location of QTL identified in previous studies of HMS.

## Discussion

### Genetic architecture of HMS

Our results support a growing body of evidence for HMS modifier alleles in mice. Several QTL have been mapped in both laboratory crosses and wild-caught mice (White *et al.* 2011; Turner and Harr 2014; Turner *et al.* 2014; Larson *et al.* 2018). However, the ability to identify these modifiers has been hampered in most previous studies by segregating variation at the major HMS loci. All the MD hybrid males in this study shared the *Prdm9^Dom2/Msc^* genotype, identical Y chromosomes (Yang (Yang *et al.* 2012; Morgan *et al.* 2016), and the PWK Chr X. Nonetheless, among these hybrids there were three distinct fertility trajectories. These distinct fertility profiles require at least two HMS modifier alleles that segregate among the four classical inbred mouse strains in our study.

PWKDBA2 hybrid males were fertile throughout life. This demonstrated that the heterozygous *Prdm9^Dom2/Msc^* genotype in combination with a PWK Chr X is not sufficient to induce HMS-related phenotypes in all genetic backgrounds. All previous studies of hybrid males carrying this Chr X and *Prdm9* genotype combination have reported either infertility or extreme subfertility (Flachs *et al.* 2012, 2014). Therefore, we conclude that at least one novel modifier allele harbored by the DBA2 strain rescued fertility, in spite of the presence of the two major HMS loci. This result compels us to reevaluate the hypothesis that mouse HMS is largely caused by asymmetric DSBs at recombination hotspots throughout the genome resulting from allele-specific PRDM9 binding. Under this hypothesis, the fertility of PWKDBA2 hybrids could be explained by a higher similarity between the PWK and DBA2 genomes relative to the other strains. However, our analysis showed that *M. m. musculus* ancestry contributed only a small fraction to the classical inbred strain genomes, and this fraction was roughly equal among the strains. We conclude that specific HMS modifier alleles are a more likely explanation. These modifiers may interact directly or indirectly with PRDM9 binding, or act independently of PRDM9. Since even infertile PWK129S1 hybrids escape complete meiotic arrest, the modifiers likely act downstream of *Prdm9* during late spermatogenesis. On the other hand, the contrasts between MD and reciprocal DM hybrids were consistent with the PWK Chr X being a necessary factor for HMS. Even the fertile PWKDBA2 hybrid had lower sperm counts than its reciprocal DBA2PWK hybrids, suggesting a pervasive negative effect of the PWK Chr X.

### Mechanisms of age-dependent HMS

Clear differences among the reproductive parameters and fertility trajectories of the hybrids suggest some HMS modifiers have age-dependent modes of action. However, the fact that there were age-dependent changes in sperm count in nearly all the hybrid types examined here suggests that hybrid male reproductive fertility is shaped by complex interactions of multiple segregating HMS alleles and age.

Histological analyses suggested that meiosis was disrupted or inefficient in infertile hybrid males by age eight weeks, as indicated by a depletion of round spermatids. This is consistent with a robust literature describing meiotic mechanisms for HMS (Good *et al.* 2008b; Mihola *et al.* 2009; Bhattacharyya *et al.* 2013; Schwahn *et al.* 2018). However, none of the MD hybrids in this study exhibited as severe meiotic impairments as seen in examples of mouse HMS with radically disorganized testes and complete arrest of meiotic prophase at or just before the pachytene substage (Flachs *et al.* 2012, 2014; Bhattacharyya *et al.* 2014). The lack of any profound meiotic disturbance in the hybrids studied here is important in light of our observations of age-dependent fertility in PWKB6 and PWKAJ. The subsequent onset of fertility later in life suggests that the eight-week old PWKB6 and PWKAJ males were just below the minimum required sperm count for fertility, and subtle phenotypic changes were sufficient to elevate a fraction of hybrid males to the fertility threshold at age 20 weeks.

Initially, we anticipated that this fertility would be accompanied by an apparent mitigation of meiotic inefficiency, manifested by an increased PCT_ROUND. However, we observed only a modest increase in PCT_ROUND in PWKB6 and PWKAJ hybrids. Instead, transient fertility was accompanied by increased productivity that appeared to be post-meiotic in nature. The most striking phenotypic differences between eight week old and 20 week old PWKB6 hybrid males was an increase in the ratio of PCT_CONDENSING to PCT_ROUND, which meant that more of the germ cells that successfully completed meiosis successfully formed mature spermatozoa. We conclude that increased post-meiotic success can increase the likelihood of fertility between the ages 12-35 weeks in PWKB6 and PWKAJ. Furthermore, this capacity is controlled by segregating genetic variation, since PWK129S1 hybrids (also MD) show no increase in post-meiotic capacity with age, and no age-dependent fertility. The four DM hybrids and PWKDBA2 also showed changes in reproductive parameters with age. However, we could not observe a similar increase in post-meiotic capacity in the five fertile hybrids, because both cell types were present in nearly all seminiferous tubules. This may reflect the limitations of our methods. Individual cell counts might reveal a similar pattern in the fertile hybrids, but quantifying this would require automated image analysis that was beyond the scope of this study, or new experiments designed to directly measure the abundance of each cell type.

The developmental origins of this age-dependent increase in post-meiotic capacity remain unknown, and we can only speculate about possible biological mechanisms. For example, testosterone levels could change with age and impact fertility (Singh *et al.* 1995; Zirkin and Tenover 2012; Beattie *et al.* 2015; Wang *et al.* 2017). Seminal vesicle weights are correlated with serum testosterone and increased with age (p ≤ 0.001) in our mice, though seminal vesicles were heaviest in infertile 35-week old hybrid males. Mitochondrial causes of age-related effects can be eliminated since all the MD hybrids inherited their mitochondria from a PWK dam, although mitochondrial-autosomal interactions cannot be ruled out. Age-related epigenetic modifications in gene regulation may also contribute to the variation we have observed. Aberrant gene expression of Chr X has been repeatedly associated with HMS (Turner and Harr 2014; Turner *et al.* 2014; Larson *et al.* 2017) and a compelling hypothesis is that HMS alleles impair both meiotic sex chromosome inactivation (MSCI) and postmeiotic sex chromatin repression (PSCR) (Bhattacharyya *et al.* 2013; Campbell *et al.* 2013). PRDM9 has a well-characterized epigenetic role in germ cell development (Parvanov *et al.* 2010; Baudat *et al.* 2010b; Baker *et al.* 2014, 2015; Davies *et al.* 2016), and it remains an intriguing question as to whether the HMS modifier alleles segregating in this study interact directly or indirectly with PRDM9.

### Implications of age-dependent HMS

Our results have immediate application to the increasingly popular Collaborative Cross and Diversity Outbred multi-parent mouse reference populations. The CC is a panel of recombinant inbred lines that are descended from eight inbred mouse strains including PWK, B6, A/J, and 129S1 (Churchill *et al.* 2004; Chesler *et al.* 2008; Aylor *et al.* 2011), and the DO is an outbred population descended from the same progenitors (Churchill *et al.* 2012). The success of the CC breeding design necessarily depended on the MD hybrids derived from those strains. PWK129S1 hybrids were found to be infertile early in the breeding program (Chesler *et al.* 2008) and did not contribute to CC lines. There was no specific observation of sterility in PWKAJ, PWKB6, or other hybrids offspring of PWK dams. However, most incipient CC lines (95%) stopped producing offspring during the inbreeding phase of the breeding program and were declared extinct. Over half of these extinct CC strains were found to display male infertility (Shorter *et al.* 2017), implicating segregating HMS alleles among all founders of the CC, not just 129S1.

The dynamic nature of fertility in PWKB6 and PWKAJ hybrid males could be critical in explaining their contributions to the CC breeding program. We conclude that a subset PWKB6 and PWKAJ males could have contributed to the CC breeding program during a narrow window of life as a result of incompletely penetrant, age-dependent fertility. Furthermore, these results could have implications for current and future studies in the CC. The facility core that provides CC mice (Welsh and McMillan 2012) reported that fewer than 50% of males sire litters for several CC strains, suggesting that HMS modifier alleles continue to segregate in the lines. Our results suggest that age may be an important design factor for CC studies that involve breeding males.

In the wild, steep allelic clines (Turner *et al.* 2012; Turner and Harr 2014) and reduced introgression on Chr X (Payseur *et al.* 2004; Janousek *et al.* 2012) provide evidence that segregating HMS alleles are a partial reproductive barrier in the Central European mouse hybrid zone. Processes that attenuate the restriction of gene flow between these subspecies are the relative permissibility of *M. m. domesticus* Chromosome X introgression due to asymmetric HMS (Turelli and Moyle 2007; Good *et al.* 2008b) and relatively infrequent hybrid female sterility (but see (Suzuki and Nachman 2015)). The modes of action of age-dependent HMS alleles have yet to be uncovered or characterized in the wild. If recovered from wild populations, these genetic variants would represent an important source of variation, allowing certain hybrid males to escape infertility during a narrow window of life. Males exhibiting age-dependent HMS would, on the one hand, have reduced relative fitness compared to males lacking sterility alleles. However, these males could also hypothetically propagate HMS alleles to the next generation. Understanding the identity, frequency, and spatial distribution of age-dependent HMS alleles will be required to fully characterize the fitness effects and evolutionary stability of this pattern of fertility.

Our observation of transient fertility in PWKB6 and PWKAJ hybrid males is the first evidence of age-dependent, segregating genetic modifiers of HMS in mice. Identification of these modifiers will undoubtedly contribute to a growing body of work characterizing the genetic architecture of mouse HMS. Unveiling the mechanisms of these age-dependent alleles not only will advance the study of reproductive barriers between subspecies, but also could provide new insights on male reproductive biology and spermatogenesis generally. These hybrids have the potential to reveal the complex interdependence of genetic variation, the aging process, and mammalian fertility.

## Acknowledgements

We thank Nicole Allard, Thomas Konneker, Erika Propst, Caitlin Varner, Connor McKenney, and Pei-Li Yao for helpful feedback and technical assistance; Tanmoy Bhattacharyya for critical input and technical assistance in histological analysis; and Fernando Pardo-Manuel de Villena, Jim Crowley, and Tim Bell for support of our initial breeding experiment. We also thank anonymous reviewers for thoughtful and helpful suggestions that improved our manuscript.

## Supporting Information Legends

**Table S1. Phenotype summary statistics.** In each hybrid male, we measured combined testes weight, sperm count, percentage of seminiferous tubules containing round spermatids (PCT_ROUND), and the percentage of seminiferous tubules containing condensing spermatids (PCT_CONDENSING) (Mean ± SE).

**Table S2. Genomic regions exhibiting specific patterns of subspecies haplotype sharing.** We identified four patterns of subspecific haplotype sharing that match the three fertility trajectories in our results. For example, the pattern in which PWK and DBA2 share a M. m. musculus origin and AJ, B6, and 129S1 share a M. m. domesticus origin is represented by the nomenclature [PWK, DBA2 | 129S1, AJ, B6]. Chromosome, start sites, stop sites, and interval width are expressed in megabases (Mb).

**Table S3. Phenotype data.** A.csv file including all phenotype data for 225 male mice (94 DM, 131 MD) that were bred at NC State. Fertility status was measured for 72 mice (17 DM, 55 MD).

**Table S4. Fertility testing to establish the cessation of fertility.** A separate experiment was conducted at UNC-CH to establish the maximum age of fertility in PWKB6 mice. All mice were infertile by age 35 weeks.

**Figure S1. Testes weights scaled by body weight**. DM hybrid males (red borders) had consistently heavier testes than MD hybrid males (black borders) when scaling by body weight, with the exception of the fertile DBA2-derived hybrids (light blue). This is the same pattern seen when not taking body weight into account (Figure 2).

